# Atom-level backbone engineering preserves peptide function while enhancing stability

**DOI:** 10.1101/2025.10.06.678421

**Authors:** Mingzhu He, Kai Fan Cheng, Anh Vu, Marcelo D. T. Torres, Stavros Zanos, Ibrahim T Mughrabi, Cesar de la Fuente-Nunez, Yousef Al-Abed

## Abstract

Peptide therapeutics offer unmatched potency and selectivity but are limited by rapid proteolysis and poor pharmacokinetics. Backbone engineering provides a rational approach to enhance stability while preserving function, yet direct comparisons across strategies remain scarce. Using bradykinin as a model, we systematically evaluated four backbone modifications—D-amino acid substitution, N-methylation, α-methylation, and azapeptide incorporation. Each modification produced distinct outcomes in synthesis, conformation, proteolytic stability, and receptor pharmacology. While D- and N-methyl substitutions yielded high stability, they compromised receptor binding and *in vivo* function. In contrast, the azapeptide analogue maintained native-like affinity and physiological activity while achieving an enhanced stability profile. These findings highlight the need to balance stability and function in peptide design and position azapeptides as an underexplored class with strong therapeutic potential. More broadly, this study establishes a framework for systematic, data-driven peptide design and optimization.

## Introduction

Peptide therapeutics have emerged as an important class of medicines that bridge the gap between small molecules and biologics. Over the past two decades, their clinical use has expanded steadily across metabolic disorders, oncology, infectious diseases, and rare conditions (1). This growth reflects their specificity, potency, and relatively favorable safety profiles. The trend is expected to accelerate, as pharmaceutical pipelines increasingly embrace peptides as efficient and versatile therapeutic agents (2, 3). Despite these advantages, native peptides are rapidly degraded by proteases and cleared by renal filtration, which restrict plasma half-lives to only minutes (1). Without chemical modification, most peptides are unsuitable for chronic therapy. Overcoming these barriers requires strategies that stabilize the peptide scaffold while retaining or enhancing biological activity.

Backbone engineering— atomic-level modification of the peptide scaffold —offers a powerful means to improve stability, with strategies such as stereochemistry changes, alkyl group additions, backbone extension or shifting and incorporation of non-natural residues (4). However, most applications remain heuristic, with single modifications targeting presumed proteolytic hotspots. While many of these efforts extend half-life, few studies systematically compare strategies across broader performance metrics, including synthetic yield, conformational dynamics, proteolytic stability, receptor binding, and *in vivo* efficacy. As a result, backbone engineering has advanced more by trial and error than by rational data-driven design, underscoring the need for frameworks that integrate multiple parameters into peptide optimization.

Here, we present a rational framework for backbone engineering (Figure 1). Using as a model bradykinin (BK), a peptide that plays a crucial role in inflammation, pain, and vasodilation, we compare four backbone modifications—D-amino acid substitution, N-methylation, α-methylation, and azapeptide incorporation—across synthetic, structural, stability, and functional parameters. Particular attention is given to azapeptides, a class of peptidomimetics with strong potential but limited adoption due to past synthetic challenges.

**Figure 1.**
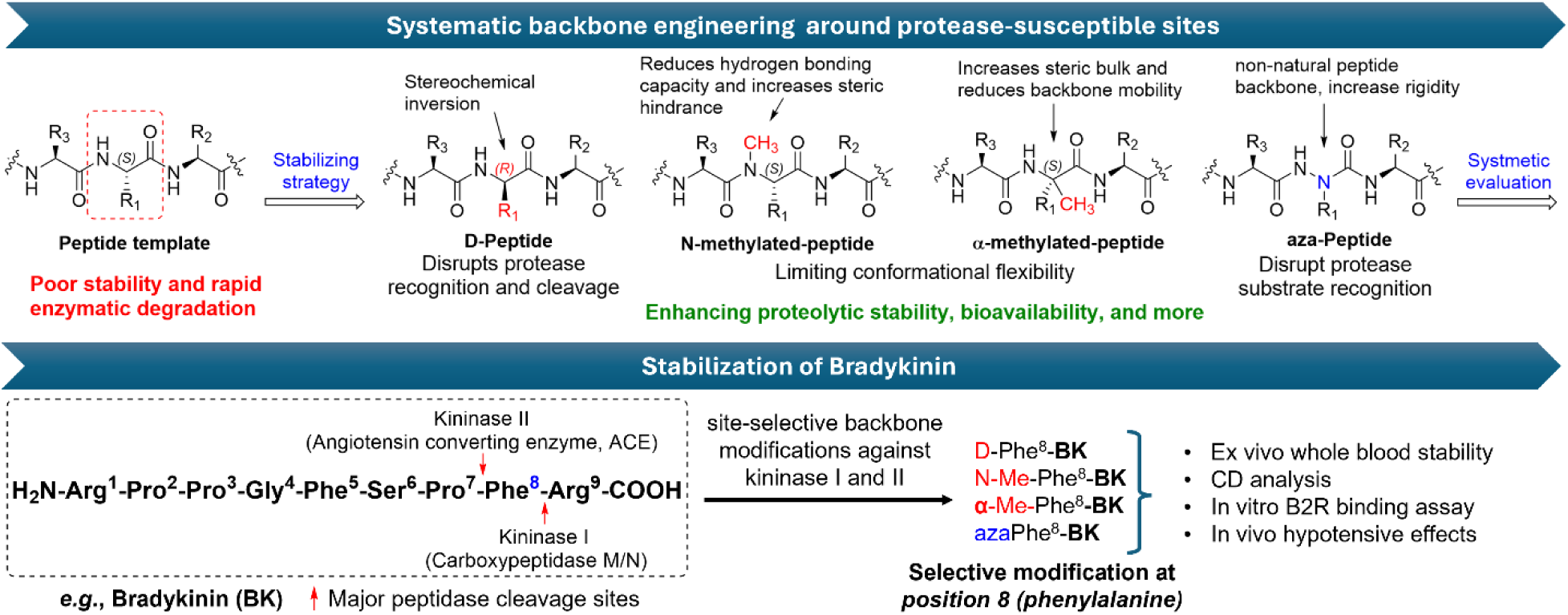
Framework for site-selective backbone engineering of bradykinin. Overview of a systematic framework to evaluate backbone modifications across synthetic, structural, stability, and functional parameters. Bradykinin (BK) was chosen as a model and modified at the Phe^8^ hotspot, a key site for kininase I and II cleavage. Four backbone modifications—D-amino acid substitution, N-methylation, α-methylation, and azapeptide incorporation—were introduced to compare their effects on *ex vivo, in vitro* and *in vivo* activity.

## Results and Discussion

Bradykinin, a nonapeptide mediator of vasodilation and inflammatory signaling, exerts its effects primarily through the B2 receptor (5, 6). BK is an ideal model for stabilization studies because its catabolic pathways are well characterized: it is rapidly degraded *in vivo* by proteases such as angiotensin-converting enzyme (ACE), neprilysin, and carboxypeptidases, with a dominant route of inactivation occurring through cleavage at the C-terminal Phe^8^ residue by carboxypeptidase N (7). This makes BK particularly suitable for evaluating backbone engineering strategies targeting proteolytic hotspots.

We designed and generated four bradykinin (BK) analogues by site-selective modification of Phe^8^ with a distinct backbone engineering strategy: D-amino acid substitution, N-methylation, α-methylation, or aza-amino acid incorporation (Figure 2). All peptides were synthesized by microwave-assisted SPPS under standardized conditions to enable direct comparison across modifications. Most analogues were obtained within one hour with yields of 45-90%. N-methylation reduced yields, likely due to cis/trans isomerization. Synthesis of the aza-analogue required extended time in accordance with our optimized protocol (8), which includes aza-amino acid integration and post-elongation (Figure 2A).

**Figure 2.**
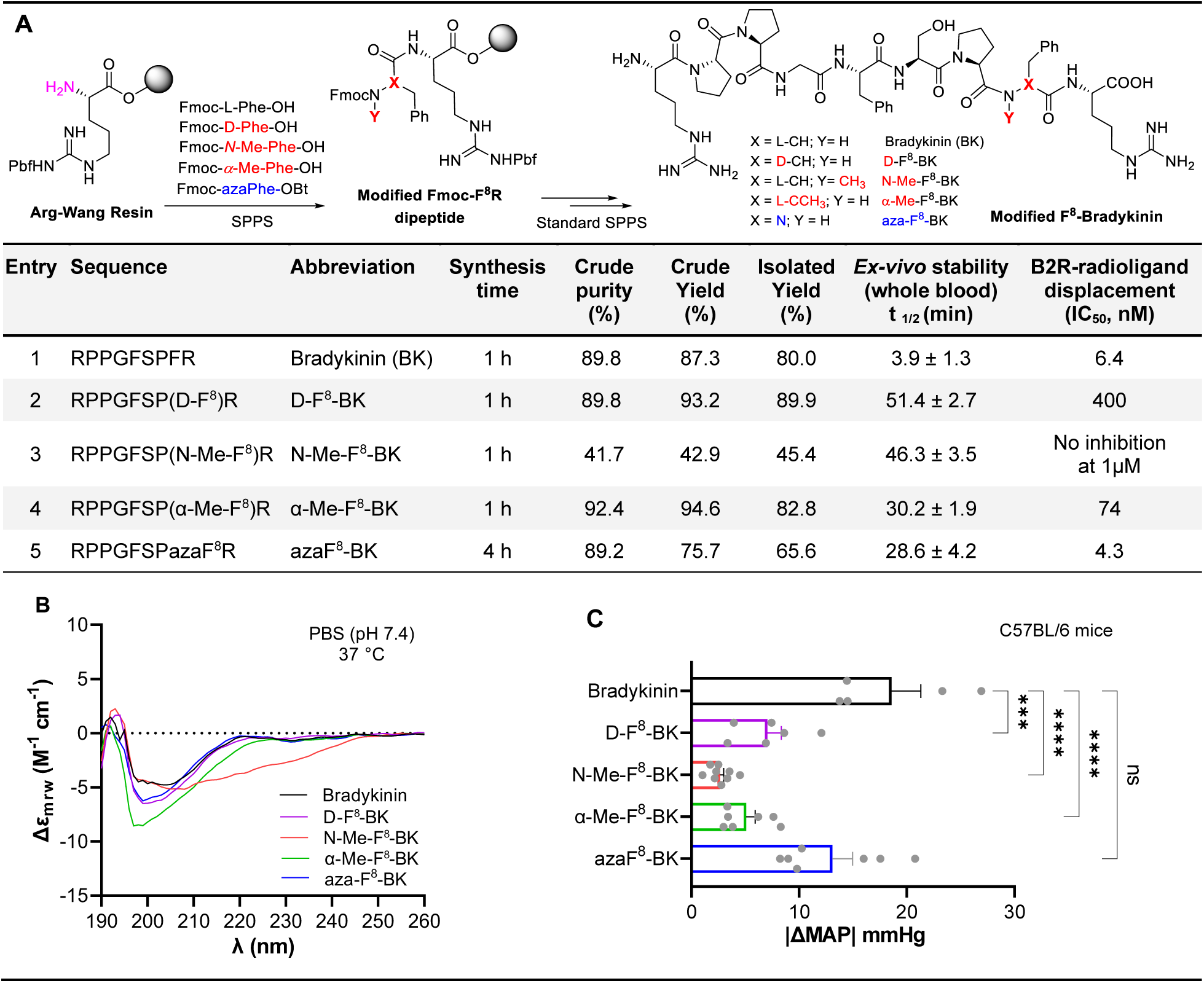
Comparative analysis of backbone modifications at Phe^8^ in bradykinin. **(A)** Standard SPPS reactions for generating bradykinin-based peptides, each modified at the Phe^8^ position with D-amino acid substitution, N-methylation, α-methylation, or aza-amino acid incorporation. All were synthesized by microwave-assisted SPPS under standardized conditions with respectable yield, crude purity of each peptide was analyzed by HPLC with the detection at 215 nm. *Ex-vivo* stability and half-lives of bradykinin-based peptides in male C57BL/6 mouse whole blood were analyzed by LCMS/MS (means ± SD, n = 3). Receptor binding (human B2R) activities (IC_50_’s) of bradykinin analogues were assessed using standardized agonist radioligand displacement assays. **(B)** Circular Dichroism (CD) spectra of bradykinin-based peptides. Bradykinin analogues were scanned from 190 to 260 nm on CD spectrometer; the spectra were plotted by using the average value of three repeats. **(C)** Hemodynamic effects (hypotensive) of intravenous bolus injections (50 μg/kg) of bradykinin and bradykinin analogues indicated by absolute change in mean arterial pressure (MAP) in C57BL/6 mice. Each grey data point represents one animal, n = 5-9. Data are presented as mean ± SEM and analyzed by one-way ANOVA with Tukey’s multiple comparisons test (bradykinin vs. test article groups): ns, not significant, ***p < 0.001, ****p < 0.0001.

All modified peptides displayed enhanced *ex vivo* stability relative to native BK (t_1/2_= 3.9 ± 1.3 min in mouse whole blood). D-F^8^-BK and N-Me-F^8^-BK showed the greatest resistance (t_1/2_= 51.4 ± 2.7 and 46.3 ± 3.5 min, respectively), followed by α-Me-F^8^-BK (30.2 ± 1.9 min) and aza-F^8^-BK (28.6 ± 4.2 min), which demonstrated moderate improvements. Circular dichroism spectroscopy revealed distinct conformational effects (Figure 2B): N- and α-methylation restricted backbone flexibility, D-amino acid substitution caused minimal deviation, and the aza-analogue preserved conformational adaptability. B2R radioligand displacement assay (9, 10) highlighted functional trade-offs. Only aza-F^8^-BK retained nanomolar affinity (IC_50_ = 4.3 nM), comparable to native BK (6.4 nM). The α-methyl analogue showed moderate binding (74 nM), whereas D- and N-methyl substitutions exhibited poor receptor interaction. *In vivo* vascular reactivity (hypotensive) assays (10, 11) mirrored these findings (Figure 2C): aza-F^8^-BK preserved receptor-mediated physiological activity, while other modifications compromised function despite improved stability.

Together, these data underscore a key principle: backbone stabilization must be balanced with functional preservation. Our bradykinin case study demonstrates the value of systematic comparisons across backbone modifications, uncovering trade-offs that single-mode evaluations would miss. Among the strategies tested, azapeptides stand out as a promising class, achieving moderate stability gains with preserved receptor pharmacology and *in vivo* activity.

Looking forward, peptide design can increasingly shift from trial-and-error approaches to predictive, experimentally validated frameworks. Establishing comparative datasets across diverse peptide targets will allow medicinal chemists to select modifications tailored to each therapeutic context, accelerating the development of stable, effective, and broadly accessible peptide drugs.

## Materials and Methods

Detailed synthetic, analytical, *ex vivo, in vitro*, as well as *in vivo* materials and methods can be found in Supplementary Information. All animal procedures were approved by the Feinstein Institutes for Medical Research Institutional Animal Care and Use Committee (IACUC, protocols #24-0260, #25-0403) and adhere to current guidelines.

## Acknowledgments

We would like to thank Dr. Barney Yoo from the Mass Spectrometry facility at Hunter College for acquiring HRMS data. This work was supported by funding from the Feinstein Institutes for Medical Research. Some of this work was supported by Northwell Health’s 2019 Innovation Challenge prize. Research reported in this preprint was supported by the National Institute of General Medical Sciences of the National Institutes of Health under award number R35GM138201 and by the Defense Threat Reduction Agency under award number HDTRA1-21-1-0014. The content is solely the responsibility of the authors and does not necessarily represent the official views of the National Institutes of Health or the Defense Threat Reduction Agency.

This preprint has not been peer-reviewed and should not be used to guide clinical.

## Author Contributions

M.H., K.F.C., and Y.A. designed the study; M.H., K.F.C., A.V., M.D.T.T, and I.T.M., ran experiments and analyzed all data; and Y.A. wrote the initial draft of the manuscript; and all authors contributed to manuscript revision and editing.

## Competing Interest Statement

K.F.C., Y.A. are on a patent application held by the Feinstein Institutes for Medical Research (FIMR) related to the azapeptide synthesis platform-Preparation of O-benzotriazole and O-imidazole synthons for use in the synthesis of peptidomimetics including azapeptides, WO2020227594 A1 2020-12-11 (active). Y.A. is inventor (FIMR) on Synthesis and uses of peptidomimetics including azapeptides, WO2020227588 A1 2020-11-12 (active). C.F.-N. is a co-founder and scientific advisor to Peptaris, Inc., provides consulting services to Invaio Sciences and is a member of the Scientific Advisory Boards of Nowture S.L., Peptidus, and Phare Bio. C.F.-N. is also on the Advisory Board of the Peptide Drug Hunting Consortium (PDHC). M.D.T.T. is a co-founder and scientific advisor to Peptaris, Inc. All the other authors declare no competing interests.

## Notes

### Summary of Updates

This revision has been submitted to (1) update the copyright license to CC BY-NC-ND (Attribution - NonCommercial - NoDerivatives), (2) update the NIH funding information, and (3) clearly state that this preprint has not been peer-reviewed. No changes have been made to the scientific content, figures, or data presented in the original submission.

