## Supplemental information for "Atom-level backbone engineering preserves peptide function while enhancing stability"

### Supplementary Information

#### 1. CHEMISTRY

**1.1 Chemicals**
Preloaded Fmoc-L-Arg(Pbf) Wang resin (100-200 mesh, 0.25 mmol/g), and Fmoc amino acids were purchased from CEM Corporation, Matthews, NC. All solvents, and reagents used in synthesis are peptide-grade and were purchased from Gyros Protein Technologies or from Sigma-Aldrich (St. Louis, MO). The solvents and reagents were used without further treatment or drying.

**1.2 Instrumentation for synthesis, purification and chemical characterization**
Fully automated solid phase peptide synthesis was performed on a Liberty Blue 2.0 Automated Microwave Peptide Synthesizer (CEM Corporation, Matthews, NC) using Preloaded Wang resin or Rink Amide Resin at 0.1 mmol scale through Fmoc-chemistry. Liberty Blue 2.0 features a unique one-pot coupling and deprotection chemistry at elevated temperature. The aza-amino acids building blocks were synthesized in our labs (ref) and automatically integrated in the solid phase peptide synthesis. Analyses were performed using Water's HPLC (Breeze Qs and Breeze 2) equipped with a 1525 binary pump, and the use of analytical column (Phenomenex kinetex 2.6 mm EVO C18 analytical column 100Å 150x4.6 mm). Chromatography was performed at ambient temperature with flow rate of 1.0 mL/min with linear gradient from Water (0.05% TFA): $CH_3CN$ (0.05% TFA) [95:5] to Water (0.05% TFA): $CH_3CN$ (0.05% TFA) [5:95] in 15 minutes and resolved peaks were detected by 2998 photodiode Array (PDA) Detector at 254 and/or 215 nm and characterized by low resolution mass spectrometry instrument (Thermo Scientific LTQXL ™) with ESI ion-source and positive mode ionization and high resolution mass spectrometry instrument (Agilent 6550 iFunnel QTOF LC/MS). Purification of the all peptidomimetics were performed on preparative HPLC purification system (Waters Prep 150 LC system combining 2545 Binary Gradient Module using XSelect Peptide CSH C18 OBD Prep Column, 130Å, 5 µm, 19 mm X 150 mm. Chromatography was performed at ambient temperature with a flow rate of 18 mL/min with a linear gradient from

Water (0.05% TFA): $CH_3CN$ (0.05% TFA)[95:5] to Water (0.05% TFA): $CH_3CN$ (0.05% TFA) [5:95] in 12 minutes, monitored by 2998 Photodiode Array (PDA) Detector UV at 254 nm and/or 215 nm.

#### 1.3 General procedure for microwave assisted solid phase peptide synthesis (SPPS)

All solid phase peptide couplings were performed at 90°C using Liberty Blue 2.0 Microwave Peptide synthesizer from CEM Corporation following standard protocol which is described sequentially below:

1) Swelling: The resin (preloaded Wang resin, 0.1 mmol) was swelled for 5 min in DMF at room temperature.
2) Fmoc Cleavage: the protected amino acid/or peptide was mixed for 1 min with 10% piperidine solution in DMF at 90°C to remove the Fmoc group. Followed by several washing steps with DMF.
3) Amino acid coupling: 5 equivalents of the next acylating component (Fmoc-protected amino acid), 5 equivalents of OxymaPure® (coupling reagent), and 10 equivalents of DIC (coupling reagent) were used to add the next amino acid in the sequence. This step is fully automated and was run according to the software installed on the synthesizer. The coupling time was set to 2 min mixing at 90°C followed by drainage step. Step 2 and step 3 were repeated until the desired peptide sequence was achieved.
4) Cleavage from the Resin: 3.0 mL of a freshly made solution of TFA/$H_2O$/TIPS (95:2.5:2.5. v/v/v) was added at 25°C to a 0.1 mmol of resin (10-25 mL/g of resin). The mixture was shaken for 2 h at 25°C, filtered, and the remaining resin was further washed with a 0.5-1.0 mL of TFA solution. The excess TFA solution was removed under nitrogen gas. The filtrate was precipitated by adding 30 mL of ice-cold ether. Upon centrifugation, the resulting solid was dissolved in 4 mL of 1:4 solution of $CH_3CN$: $H_2O$. The resulting solution was lyophilized followed by HPLC purification.

#### 1.4 General procedure for automated integration and acylation of the aza-amino acid in SPPS(1)

Fmoc-L-Arg(Pbf) Wang resin (0.1 mmol) was swelled for 10 min in DMF (10 mL), followed by Fmoc cleavage using 10% piperidine solution in DMF at 90°C to remove the Fmoc group. Followed by washing steps with DMF (4.0 mL x 4), Fmoc-azaPhe-OBt building block (1) (0.5 mmol, 5 equiv, 0.2M) in DMF (2.5 mL) was automatically transferred into the reaction vessel. The coupling time was set to 2 h mixing at 60°C followed by drainage step. The Fmoc protected aza-Phe bound to the peptidyl chain (based on 0.1 mmol peptide) was mixed with 10% piperidine solution in DMF (4.0 mL) at 90°C to remove the Fmoc group, then the resin (0.1 mmol) was suspended in DMF (4.0 mL) with Fmoc-L-Pro-OH (5 eq, 0.5 mmol), OxymaPure (5 eq, 0.5 mmol) and DIC (10 eq, 1.0 mmol). The resulting suspension was mixed for 1 h at 60°C, followed by drainage and washes with DMF. Step 2 and step 3 in 1.3 were repeated until the desired peptide sequence was achieved.

#### 1.5 Synthesis and characterization of Bradykinin analogues on a Liberty Blue 2.0 Automated Microwave Peptide Synthesizer

Bradykinin analogues were synthesized from 0.1 mmol of Preloaded Fmoc-L-Arg(Pbf) Wang resin (100-200 mesh, 0.25 mmol/g) following the general procedure for solid phase peptide synthesis. After standard SPPS, washes and resin cleavage, bradykinin analogues were analyzed by HPLC to confirm the crude yield and crude purity. Then the resulting crude material was purified using prep C18 column and linear gradient. After lyophilization, isolated yield and final product's purity of these analogues were confirmed by analytical HPLC. Bradykinin analogues were collected and submitted to HRMS analysis.

**Synthesis of the Bradykinin (H-RPPGFSPFR-OH)**

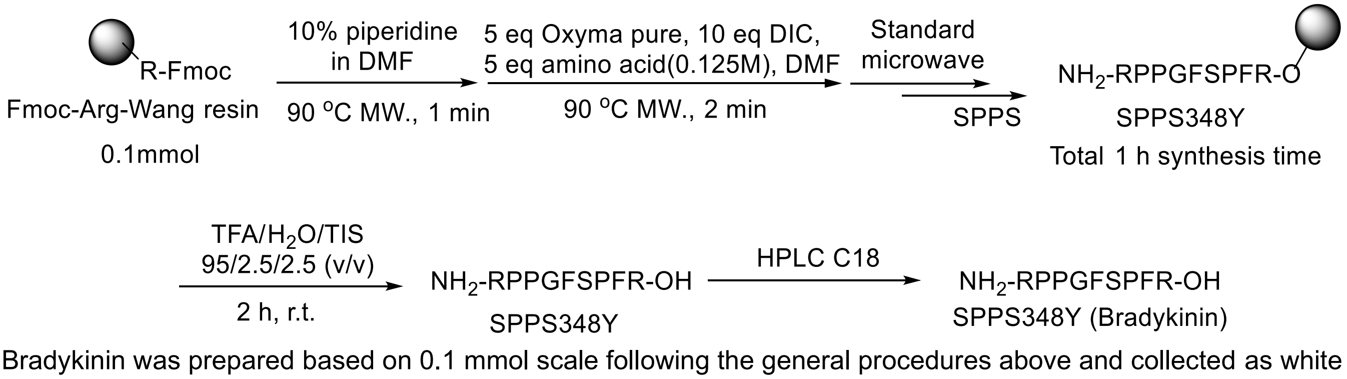


Bradykinin was prepared based on 0.1 mmol scale following the general procedures above and collected as white fluffy powder in 89.8% crude purity. The crude material was purified using prep HPLC system, after lyophilization, collected as fluffy powder (80.0% isolated yield). With purity of 98.8% based on HPLC, RT = 4.77 min (flow rate of

1.0 mL/min with linear gradient from water (0.05% TFA): $CH_3CN$ (0.05% TFA) [95:5] to water (0.05% TFA): $CH_3CN$ (0.05% TFA) [5:95] in 15 min, monitored/detected UV at 215 nM by Photodiode Array (PDA) Detector. HRMS m/z calculated for $C_{50}H_{73}N_{15}O_{11}$ [M+H] 1060.5687 found 1060.5690.

**Synthesis of the D-F$^8$-Bradykinin (H-RPPGFSP(D-F$^8$)-R-OH)**

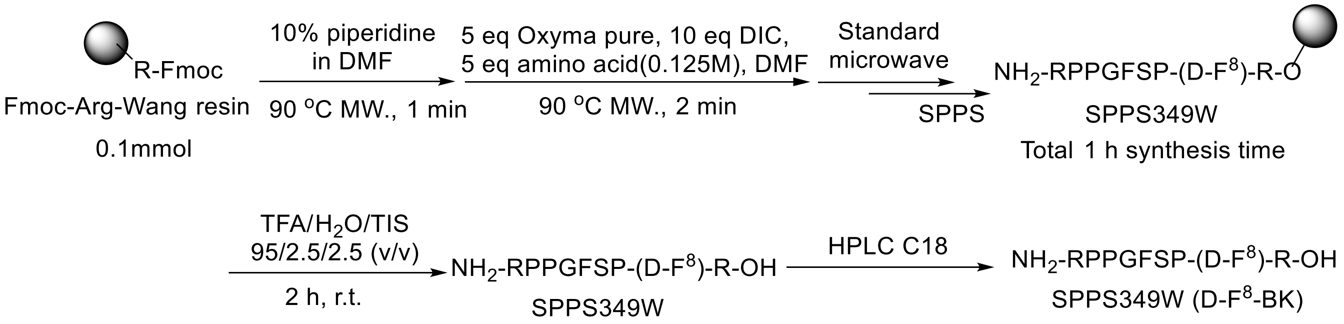


D-F$^8$-Bradykinin was prepared based on 0.1 mmol scale following the general procedures above and collected as white fluffy powder in 89.8% crude purity. The crude material was purified using prep HPLC system, after lyophilization, collected as fluffy powder (89.9% isolated yield). With purity of 98.5% based on HPLC, RT = 4.93 min (flow rate of 1.0 mL/min with linear gradient from water (0.05% TFA): $CH_3CN$ (0.05% TFA) [95:5] to water (0.05% TFA): $CH_3CN$ (0.05% TFA) [5:95] in 15 min, monitored/detected UV at 215 nM by Photodiode Array (PDA) Detector. HRMS m/z calculated for $C_{50}H_{73}N_{15}O_{11}$ [M+H] 1060.5687 found 1060.5687.

**Synthesis of the N-Me-F$^8$-Bradykinin (H-RPPGFSP(N-Me-F$^8$)-R-OH)**

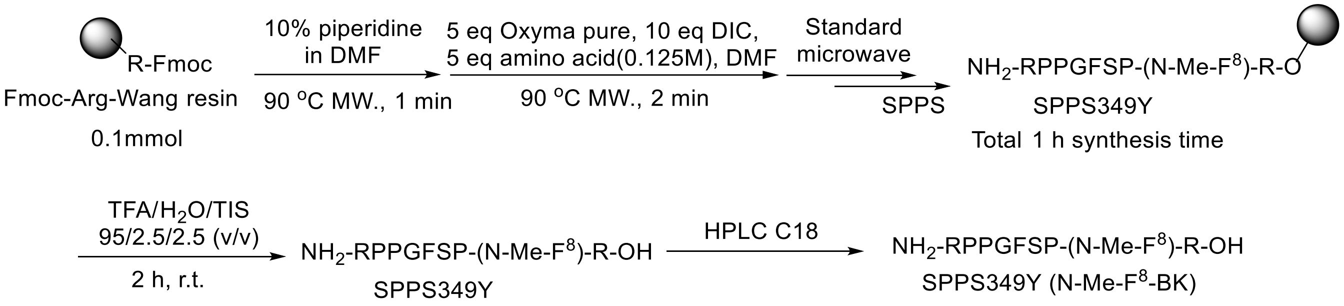


N-Me-F$^8$-Bradykinin was prepared based on 0.1 mmol scale following the general procedures above and collected as white fluffy powder in 41.7% crude purity. The crude material was purified using prep HPLC system, after lyophilization, collected as fluffy powder (45.4% isolated yield) as mixture of cis/trans isomers. With purity of 99.4% based on HPLC, RT = 5.3-5.5 min (flow rate of 1.0 mL/min with linear gradient from water (0.05% TFA): $CH_3CN$ (0.05% TFA) [95:5] to water (0.05% TFA): $CH_3CN$ (0.05% TFA) [5:95] in 15 min, monitored/detected UV at 215 nM by Photodiode Array (PDA) Detector. HRMS m/z calculated for $C_{51}H_{75}N_{15}O_{11}$ [M+H] 1074.5843 found 1074.5842.

**Synthesis of the α-Me-F$^8$-Bradykinin (H-RPPGFSP(α-Me-F$^8$)-R-OH)**

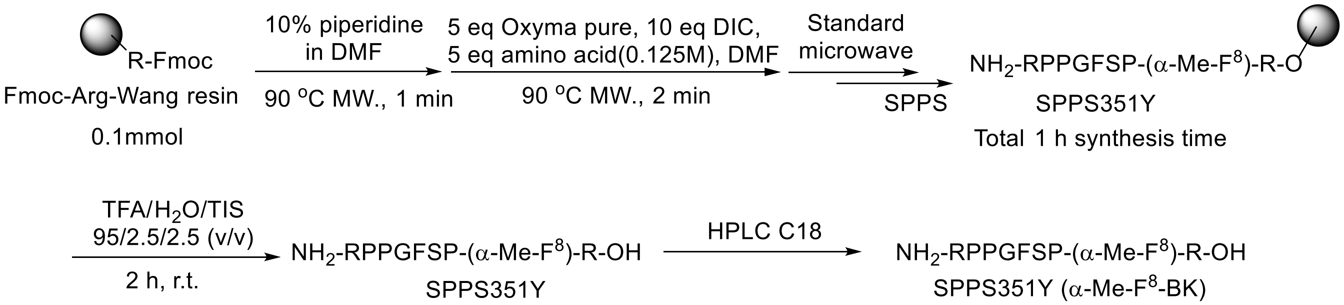


α-Me-F$^8$-Bradykinin was prepared based on 0.1 mmol scale following the general procedures above and collected as white fluffy powder in 92.4% crude purity. The crude material was purified using prep HPLC system, after lyophilization, collected as fluffy powder (82.8% isolated yield). With purity of 99.6% based on HPLC, RT = 5.1 min (flow rate of 1.0 mL/min with linear gradient from water (0.05% TFA): $CH_3CN$ (0.05% TFA) [95:5] to water (0.05% TFA): $CH_3CN$ (0.05% TFA) [5:95] in 15 min, monitored/detected UV at 215 nM by Photodiode Array (PDA) Detector. HRMS m/z calculated for $C_{51}H_{75}N_{15}O_{11}$ [M+H] 1074.5843 found 1074.5843.

**Synthesis of the aza-F$^8$-Bradykinin (H-RPPGFSPazaF$^8$-R-OH)**
azaF$^8$-Bradykinin was prepared based on 0.1 mmol scale following the general procedures above and collected as white fluffy powder in 89.2% crude purity. The crude material was purified using prep HPLC system, after lyophilization, collected as fluffy powder (65.6% isolated yield). With purity of 98.9% based on HPLC, RT = 4.82 min (flow rate of 1.0 mL/min with linear gradient from water (0.05% TFA): $CH_3CN$ (0.05% TFA) [95:5] to water (0.05% TFA): $CH_3CN$ (0.05% TFA) [5:95] in 15 min, monitored/detected UV at 215 nM by Photodiode Array (PDA) Detector. HRMS m/z calculated for $C_{49}H_{72}N_{16}O_{11}$ [M+H] 1061.5639 found 1061.5640.

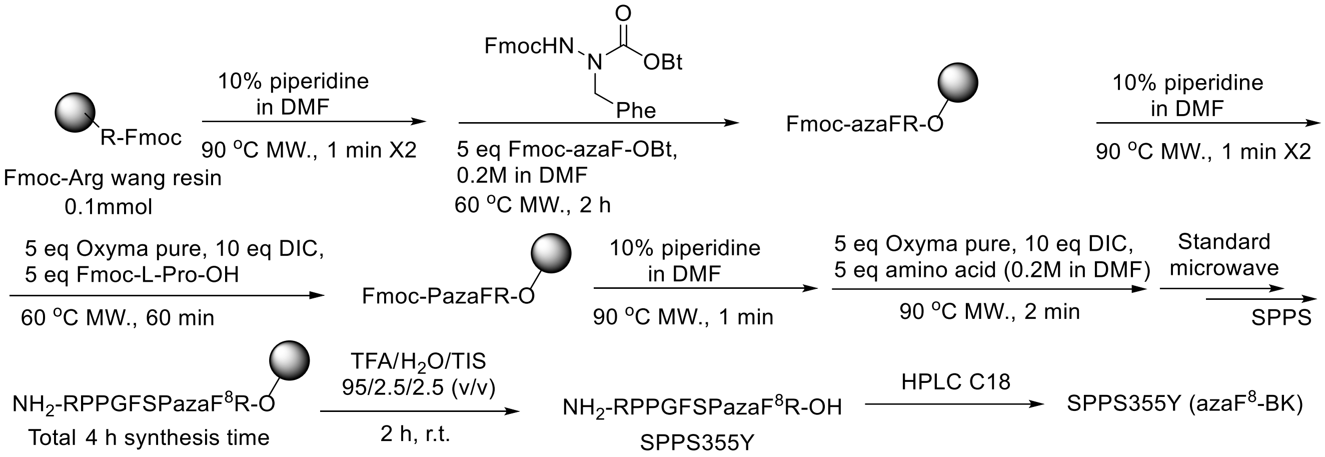


### 2. Ex vivo stabilities and half-life experiments of bradykinin-based peptides

Studies were carried out in male C57BL/6 mouse whole blood collected in sodium heparin. Blood was adjusted to pH 7.4 prior to initiating the experiments. Water stocks (5 mg/ml) were first prepared for the test article. Aliquots of the water solutions were dosed into 2 mL whole blood, which had been pre-warmed to 37°C, at a final test article concentration of 0.1mg/mL. The vials were kept in a 37°C incubator for the duration of the experiment. Aliquots (200 µ) were taken at each time point (0, 5, 10, 20, 30, 60 and 120 minutes) and added to vials which had been pre-filled with 400 µL of acetonitrile (ACN). Samples were stored at 4°C until the end of the experiment. After the final time point was sampled, the vials were mixed and then centrifuged at 18200 x g for 10 minutes. Aliquots of the supernatant were removed, diluted 1:1 into 2% ACN in 0.1% formic acid water, and analyzed by LCMS/MS. The final solution was transferred into polypropylene vials, and then 10 µl was injected and analyzed on a LTQ XL™ Linear Ion Trap mass spectrometry coupled to a Vanquish UHPLC system (Thermo Scientific, USA), with a Accucore™ Vanquish™ C18+ UHPLC Column (1.5 µm, 2.1x 50 mm) at 45ºC. A linear gradient of 5-95% acetonitrile in water (0.1% formic acid) was used for 5 min with a flow rate of 0.2 mL/min. Data were analyzed using Xcalibur 3.1 (Thermo Scientific, USA). The peak area response ratio (PARR) to calibration curve was compared to the PARR at time 0 to determine the percentage of intact peptide remaining at each time point during incubation. Half-lives were calculated using Excel. $t_{1/2}$ results are means ± SD from three blood samples, each data point was run in duplicate.

### 3. In vitro pharmacology: Human B2 (h) (agonist radioligand) receptor binding assay for bradykinin analogues

The data was collected by CRO Eurofins Cerep (France). Test compounds (BK-1 / bradykinin, BK-2 / D-F$^8$-BK, BK-3 / N-Me-F$^8$-BK, BK-4 / α-Me-F$^8$-BK and BK-5 / azaF$^8$-BK) were supplied to Eurofins in a blinded manner and run at five concentrations (0.001, 0.01, 0.03, 0.1, 1 µM) to determine the IC-50's. Standard company control test compounds were used for assay validation. Experimental assays achieved inhibition of more than 50% and were considered to represent significant effects of the test compounds. Tritium-labeled bradykinin (B2(h)) was used as the agonist for B2R receptor binding. The results are expressed as a percent of control specific binding (measured specific binding /control specific binding)*100, and as a percent inhibition of control specific binding (100 minus (measured specific binding/ control specific binding)*100) obtained in the presence of the test compounds. The IC50 values (concentration causing a half-maximal inhibition of control specific binding) and Hill coefficients (nH) were determined by non-linear regression analysis of the competition curves generated with mean replicate values using Hill equation curve fitting $Y=D+[A-D / 1+(C/C_{50})^{nH}]$ where Y = specific binding, A = left asymptote of the curve, D = right asymptote of the curve, C = compound concentration, $C_{50} = IC_{50}$, and nH = slope factor. This analysis was performed using software developed at Cerep (Hill software) and validated by comparison with data generated by the commercial software SigmaPlot® 4.0 for Windows® (© 1997 by SPSS Inc.). The inhibition constants (Ki) were calculated using the Cheng Prusoff equation. $Ki=IC_{50}/(1+L/Kd)$ where L = concentration of radioligand in the assay, and Kd = affinity of the radioligand for the receptor. A scatchard plot is used to determine the Kd.

#### 4. Circular Dichroism (CD) spectra of bradykinin-based peptides.

The circular dichroism experiments were conducted using a J1500 circular dichroism spectropolarimeter (Jasco) in the Biological Chemistry Resource Center (BCRC) at the University of Pennsylvania. Experiments were performed at 37 °C, the spectra graphed are an average of three accumulations obtained with a quartz cuvette with an optical path length of 1.0 mm, ranging from 260 to 190 nm at a rate of 50 nm min-1 and a bandwidth of 0.5 nm. The concentration of all peptides tested was 50 µM, and the measurements were performed in PBS, with baseline recorded prior to measurement. A Fourier transform filter was applied to minimize background effects. Secondary structure fraction values were calculated using the single spectra analysis tool on the server BeStSel (2).

#### 5. Surgical procedure and physiological blood pressure measurement

Animals and procedures

Adult male C57BL/6 mice (aged 8–12 weeks; n = 5–9 per group) were purchased from Charles River Laboratories (Wilmington, MA) and housed under standard laboratory conditions with ad libitum access to food and water. All animal experiments were approved by the Institutional Animal Care and Use Committee of the Feinstein Institutes for Medical Research (protocol number: 25-0403). After isoflurane anesthesia (5% induction, 1-2% maintenance), mice were placed on a heated surgical platform and instrumented with ECG leads. A 2-cm midline incision was made in the ventral neck, and the right carotid artery and right jugular vein were isolated. The jugular vein was catheterized using a 1-Fr catheter (Instech Labs, Plymouth Meeting, PA) by ligating the rostral end and momentarily occluding the caudal end to allow insertion into the vessel without bleeding. Following the same technique, the carotid artery was instrumented with a fluid-filled catheter connected to a pressure transducer (ADI Instruments, Colorado Springs, CO) for continuous blood pressure monitoring. The ECG and blood pressure signals were amplified using Bio-Amp and Bridge Amp and digitized using PowerLab 16/35 (ADI Instruments). Mean arterial blood pressure (MAP) was calculated using systolic and diastolic blood pressure measurements.

Peptide administration and blood pressure measurement

Peptides were dissolved in isotonic saline (0.9% NaCl) and injected into the jugular vein as IV boluses of 50 µl. A dose of 50 µg/kg was determined by obtaining a dose-response curve for bradykinin and choosing the dose that resulted in ~20 mmHg change in MAP, similar to previous reports (3, 4). Baseline MAP was measured for a period of 10 min before administering 3-4 boluses of a given peptide. After each bolus, a ~25 min waiting period was implemented to allow blood pressure to return to baseline before injecting a subsequent dose. At the end of the experiment, animals were euthanized by exsanguination under deep anesthesia. Delta MAP was calculated as the difference between pre-injection MAP (30 sec average immediately before injection) and post-injection MAP (30 sec average around the lowest MAP reading after injection). The first MAP measurement was not included in calculations to avoid variability caused by volume of saline in catheter tube.
